## Supplemental Figures for "Rubella virus tropism and single cell responses in human primary tissue and microglia-containing organoids"

**Figure Supplements**

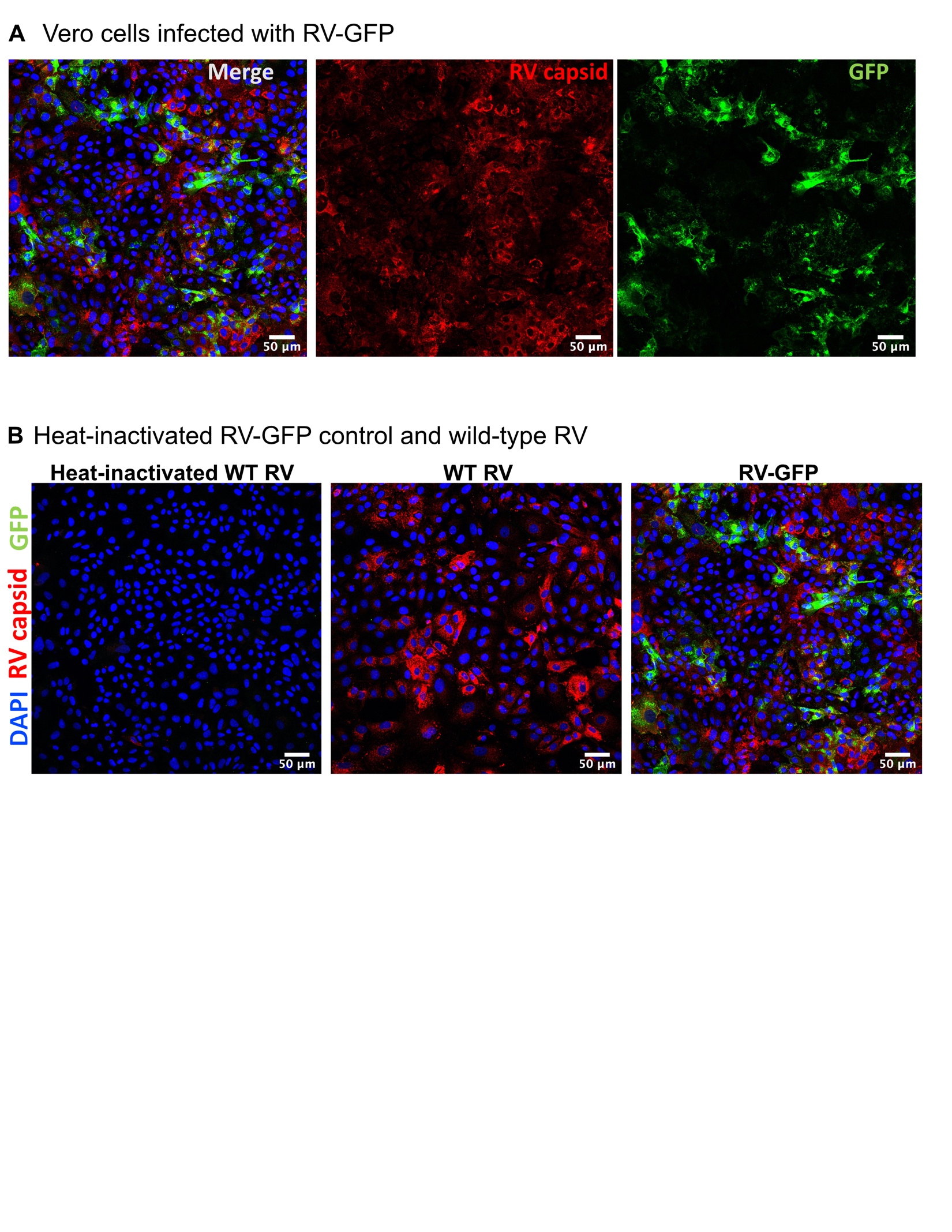

**Figure 1 – figure supplement 1. GFP expression in RV-infected Vero cells**. **A**. Co-localization of RV viral capsid and GFP expression in Vero cells. **B**. Heat-inactivated wild type RV, parental wild type RV strain and GFP-expressing RV construct in Vero cells. RV-GFP image is the same as in A.

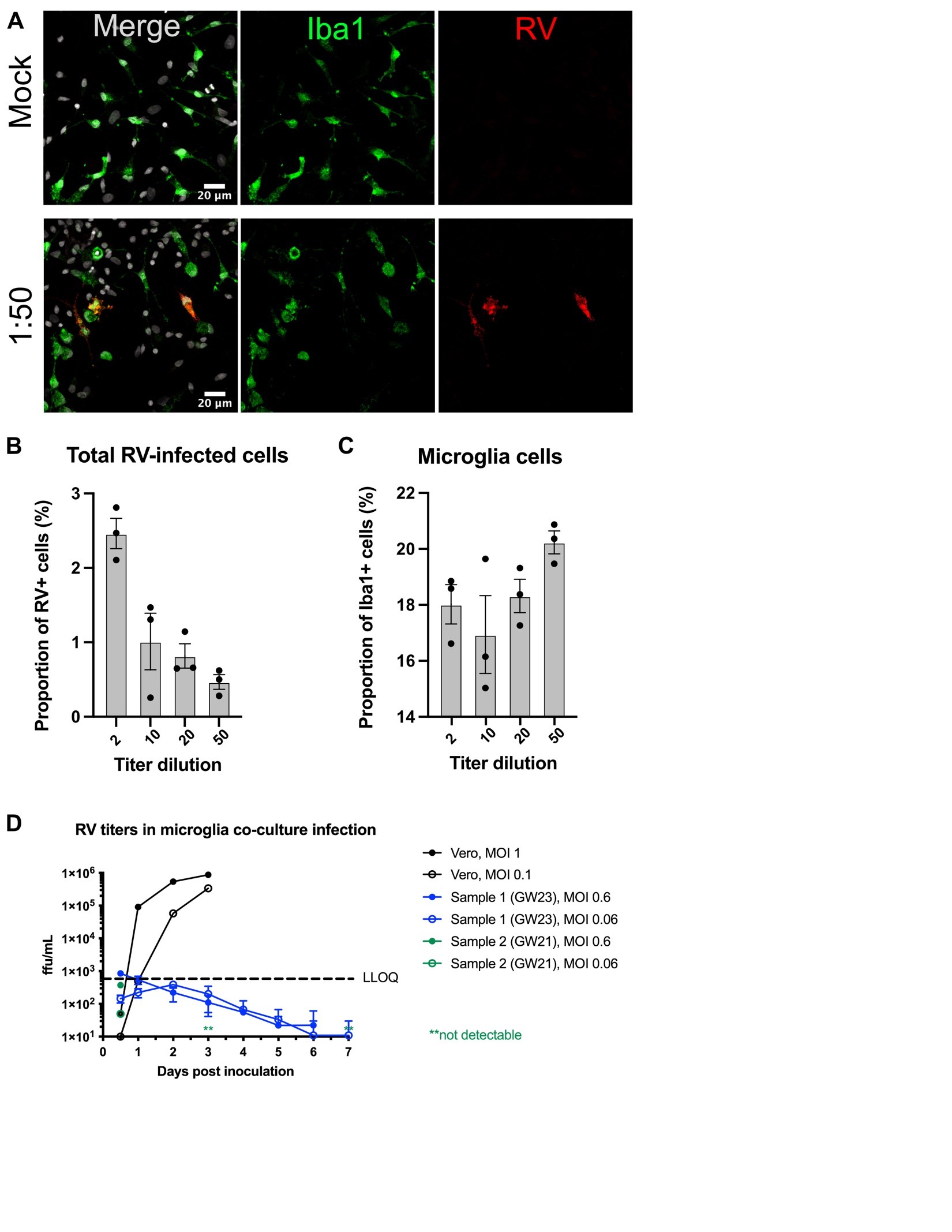

**Figure 2 – figure supplement 2. RV inoculum dilution in mixed co-cultures of microglia and non-microglia cells**. **A**. Representative images of co-cultures inoculated with mock virus or with RV viral stock diluted at 1:50. **B**. Quantifications of cells immunopositive for RV capsid across different inoculum dilutions show a decrease in the overall proportion of RV capsid-positive cells at lower titers. Columns represent mean with standard error means. Dots represent individual wells. **C**. Proportion of microglia is not changed across different viral stock dilutions. Columns represent mean with standard error means. Dots represent individual wells. **D**. Rubella virus titering experiment performed in Vero cells (positive control) or dissociated microglia co-cultures. In primary microglia co-cultures, viral titer falls below detection levels after several days of infection.

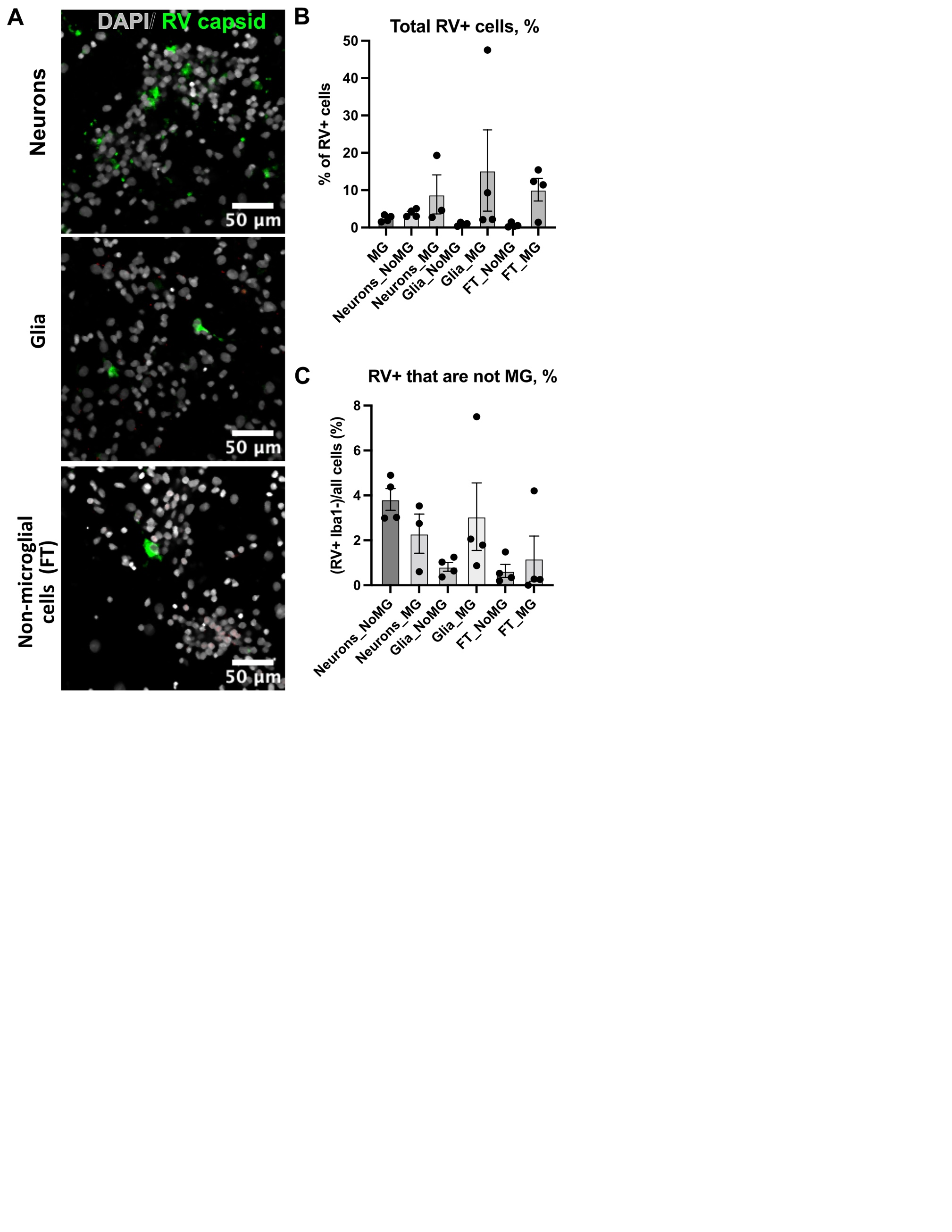

**Figure 2 – figure supplement 3. Rubella infection in non-microglia cells. A.** Representative images of different cell types depleted of microglia. Cell cultures were stained RV capsid (green) and DAPI. **B**. Quantification of total cells that are positive for RV capsid across conditions. **C**. Quantification of RV+ cells that are not microglia across different cell populations. No statistically significant difference was detected in RV infectivity in cells c-cultured with or without microglia.

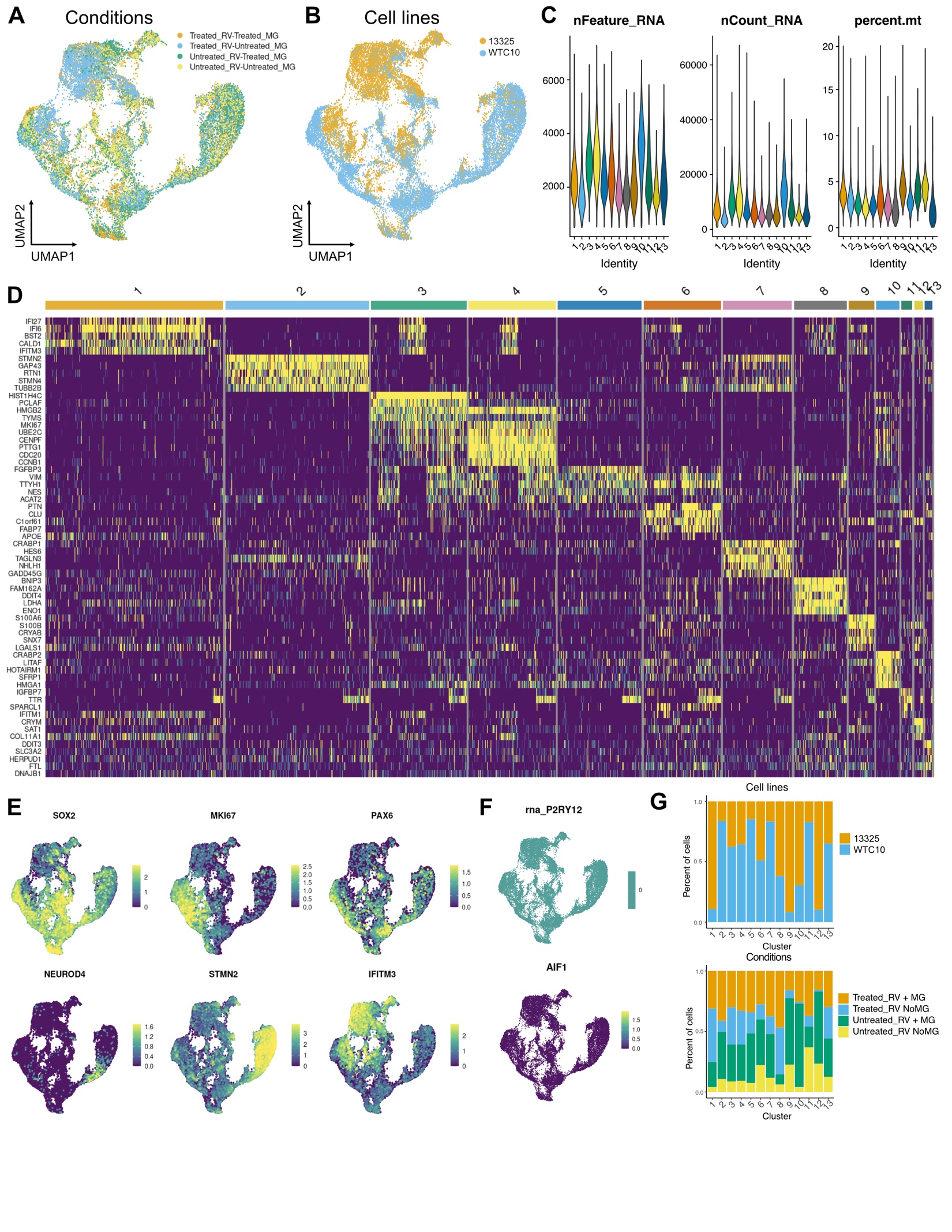

**Figure 5 – figure supplement 4. scRNAseq analysis of brain organoids**. **A**. UMAP of brain organoids colored by condition. **B**. UMAP colored by individual cell line. Two iPSC cell lines, 13325 and WTC10, were used for the scRNAseq experiment. Cell line details are described in Materials and Methods. **C**. By-cluster distribution of quality control metrics of scRNAseq data, including total number of genes detected in each cell (‘nFeature_RNA’), total number of molecules detected within a cell (‘nCount_RNA’) and percent of mitochondrial reads (‘percent mito’). **D**. Heatmap of top cluster marker genes for each cluster. **E**. Feature plots for select cluster marker genes. *SOX2* represents radial glia cells and astrocytes; *MKI67* is a marker of dividing cells; *PAX6* labels intermediate progenitor cells; *NEUROD4* is a neuronal differentiation gene and a marker for immature neurons and neural progenitor cells; *STMN2* is a marker for neurons; *IFITM3* is a marker for interferon response. **F**. Feature plots for microglia markers. Top – no canonical microglia marker *P2RY12* was detected in the dataset. Bottom – rare *AIF1*-positive cells are scattered throughout all clusters. **G**. Contribution of a cell line (top) and each condition (bottom) to each cluster composition.
